# Design and Validation of New Primers for Specific and Sensitive Real-time PCR Detection and Quantification of Seven Botulinum Encoding Genes (Serotype A-G) of *Clostridium botulinum*

**DOI:** 10.64898/2026.08.21.746353

**Authors:** Phuong-Lan Phan, Hai-Anh Chu, The-Thai Le, Phan-Anh Le, Hong-Loan Thi Nguyen, Minh-Nguyet Thi Tran, Yen Pham, Trung-Thanh Nguyen, Tuan-Nghia Phan

## Abstract

Botulinum neurotoxins (BoNTs) comprise a highly diverse group of seven serotypes (from A-G) and over 40 subtypes worldwide. Previous primer- and probe-based nucleic acid amplification tests (NAATs) for detection of BoNT encoding genes are challenged by high levels of nucleotide polymorphism both across and within subtypes. In this study, multiple BoNT gene sequences were aligned to identify highly conserved regions for the design of new primers that enable the detection of all seven serotypes under the same conditions. Specific primer sets were designed and validated using *in silico*, conventional and real-time PCR with constructed plasmids carrying the target fragments and spiked food matrices. The established procedure achieved highly specific and sensitive detection of BoNT serotypes A–G with sensitivity of 10 copies/reaction and a total turnaround time of approximately 1.5 hours. The procedure also eliminated the carryover PCR product by using uracil-N-glycosylase in combination with dUTP in the assay reaction mix. This study provides an alternative NAAT with higher coverage and compliments the traditional mouse bioassays in enhancing global botulism surveillance capabilities.

## Introduction

Botulinum (BoNT) poisoning, known as botulism, leads to flaccid paralysis by inhibiting acetylcholine release in neuronal transmission, subsequently resulting in potentially fatal respiratory failure [1,2]. Botulism occurs via several routes of exposure. Foodborne botulism results from ingestion of preformed toxin in improperly processed or preserved foods, particularly those stored under anaerobic conditions. Infant botulism, the most common form in many countries, occurs when ingested spores germinate and produce toxin in the immature intestinal tract [3]. Additional forms include wound botulism, iatrogenic botulism following therapeutic or cosmetic use of BoNT, and inhalational botulism, which is extremely rare and primarily associated with bioterrorism scenarios [2–4]. Despite low global incidence, botulism is associated with a relatively high mortality rate of approximately 5–10%, particularly when diagnosis and antitoxin treatment are delayed [5].

BoNT produced by *Clostridium botulinum* and related *Clostridium* species are categorized into seven serotypes from A to G based on immunological properties [6–9], and a proposed eighth serotype (BoNT/H), now generally regarded as a chimeric BoNT/HA toxin, requiring further characterization [10]. Human botulism is primarily caused by four serotypes A, B, E, and F, whereas C, D, and G are linked to animal botulism [11, 12]. Each BoNT serotype comprises multiple subtypes defined by amino acid sequence variation. These subtypes may differ in toxicity, receptor binding affinity, duration of action, and immunological properties, posing challenges for both diagnosis and treatment [9]. In addition, based on physiological and genomic characteristics, *C. botulinum* strains are classified into four major groups (I–IV), which differ in proteolytic capacity, optimal growth temperature, spore heat resistance, ecological niche, and toxin production profile [8]. Notably, the *bont* genes, which encode the botulinum neurotoxin, may be located on chromosomal DNA, plasmids, or bacteriophages, highlighting the importance of horizontal gene transfer in the evolution and dissemination of toxigenic clostridia [7,13]. This genetic plasticity contributes to extensive diversity in toxin structure, antigenicity, and biological activity. Hence, understanding the genetic diversity of *bont* genes is critical for epidemiological surveillance and the development of reliable diagnostic assays.

Laboratory confirmation of botulism traditionally relies on the mouse bioassay, which remains the gold standard for detecting biologically active toxin. However, this method is time-consuming, labor-intensive, and raises ethical concerns related to animal use [14]. Consequently, nucleic acid amplification techniques (NAATs), particularly polymerase chain reaction (PCR)–based assays targeting *bont* genes, have become increasingly important. Conventional PCR, multiplex PCR, loop-mediated isothermal amplification (LAMP), real-time PCR, and multiplex real-time PCR assays have been developed to detect and differentiate BoNT-producing *Clostridium* species with high sensitivity and specificity [14–19]. Although these molecular methods detect the presence of toxin genes rather than active toxin, they provide rapid and reliable alternatives for surveillance and outbreak investigations. Recently, new biosensors and whole-genome sequencing have provided complementary tools for surveillance, epidemiology, and outbreak tracing [20–23].

Despite these advances, comprehensive molecular assays capable of detecting and differentiating all seven established serotypes (A–G) face difficulties, because the botulinum neurotoxins are extremely diverse, and more *bont* gene diversity is being discovered, such diversity creates a major challenge for molecular diagnostics because sequence polymorphisms within primer-and probe-binding regions may reduce assay inclusivity and increase the risk of false-negative detection [11,19]. Given the high mortality associated with botulism and the constraints of traditional diagnostic methods, the development of rapid, sensitive, and ethically acceptable molecular detection strategies is essential. This work provides an improved real-time PCR procedure for highly specific and sensitive detection of seven serotypes (A-G) of *C. botulinum* neurotoxin genes with broad coverage of gene polymorphism. Our results are expected to facilitate early case identification, support timely clinical intervention, and strengthen food safety surveillance.

## Materials and Methods

### *C. botulinum* type A–B strains, Plasmid DNA Templates, primers pairs

Two *C. botulinum* type A–B strains were isolated from the samples that tested the positive for botulinum neurotoxin serotypes A–B (BoNT/A–B), respectively, they belong to the strain collection from food poisoning cases in 2020–2021, and were preserved at National Institute for Food control, Vietnam. They were cultured in Cooked Meat broth medium and enriched under anaerobic conditions at 37°C prior to RNA extraction. Due to biosafety regulations, it was impossible to order toxigenic *C. botulinum* strains from any culture collection. *E. coli* VTCC 12272 and *B. subtillis* VTCC 11010 were ordered from Viet Nam Type Culture Collection (Viet Nam).

Plasmids carrying selected target gene fragments corresponding to botulinum neurotoxin serotypes A–G (BoNT/A–G) were generated by either cloning the PCR-amplified genes (BoNT/A and BoNT/B) into pGEM-T vector or synthesized (BoNT/C-G) by IDT Company (The United States). These plasmids were used as DNA templates for assay development, optimization, and validation. The sequence and location of each selected BoNT gene fragments are listed in supplementary 1.

Specific primer pairs used to amplify target gene fragments for serotypes A-G were designed and ordered from PhuSa Genomic company (Vietnam). Detailed primers sequences are listed in Table 1.

**Table 1.**
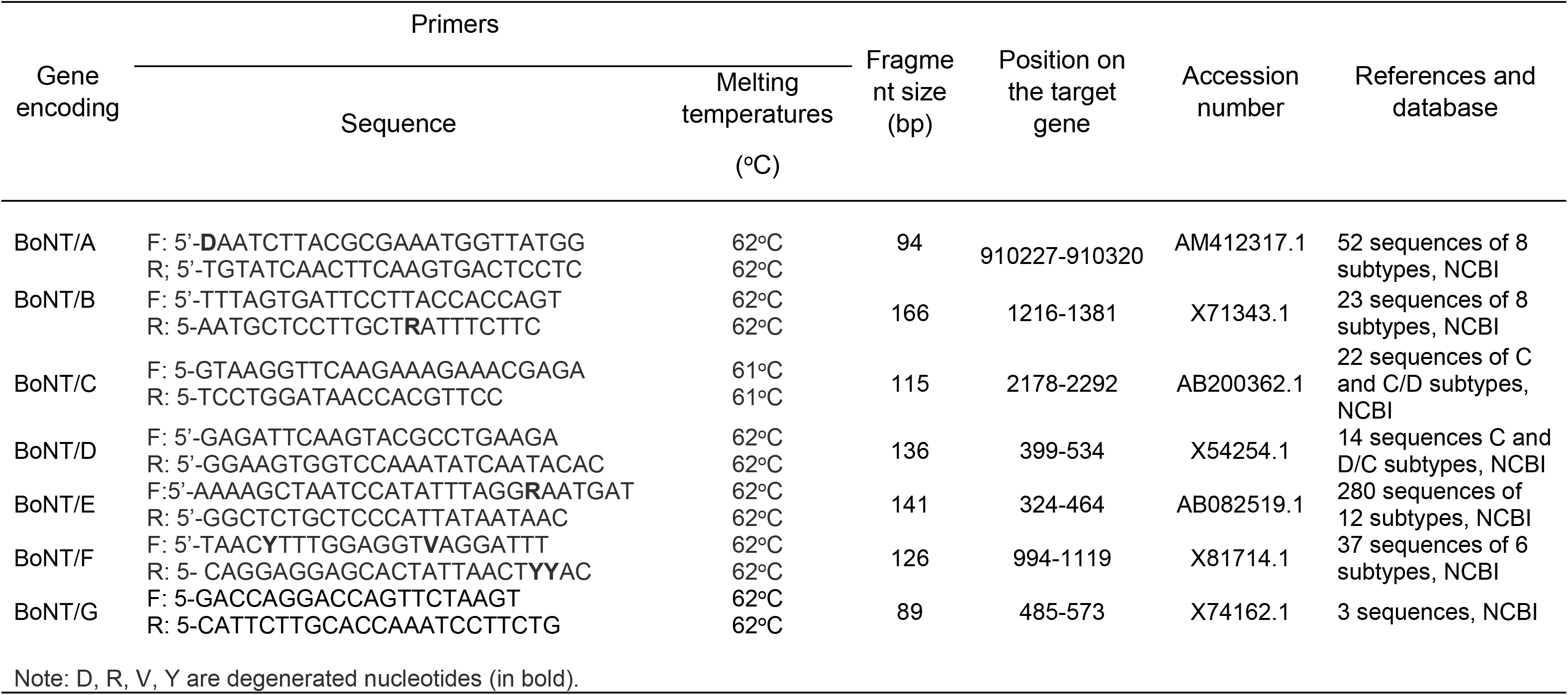
Selected conserved regions and primer sequences for the detection of BoNT/A-G encoding genes.

### Reagents and Consumables

The DNA-spin™ Plasmid DNA Purification Kit was purchased from iNtRON, Republic of Korea. PCR and real-time PCR reagents included GoTaq® Green Master Mix (Promega, United States), TaqPath™ 1-Step Multiplex Master Mix (4×; Thermo Fisher Scientific, United States), PowerUp™ SYBR Green Master Mix (2×;Thermo Fisher Scientific, United States), Luna® Universal qPCR Master Mix (2×; New England Biolabs, United States), and SYBR™ Green dye was purchased from Invitrogen (United States).

All other reagents were purchased from reputable suppliers and were of molecular biology grade.

### Equipment

PCR amplification was performed using an Eppendorf thermal cycler (Eppendorf, Germany). Real-time PCR assays were conducted using a CFX96 Opus Real-Time PCR System (Bio-Rad, United States). Additional equipment included a microcentrifuge, vertical and horizontal gel electrophoresis systems, a GelDoc imaging system (Bio-Rad, United States), and a NanoDrop spectrophotometer (Thermo Fisher Scientific, United States).

### Strain Culture

Strains of *C. botulinum* serotype A and B were grown anaerobically in deoxygenated culture broth cooked meat medium at 30°C as descried by Fohler et al. [18]. After 7 days, culture was taken out for DNA isolation.

### Preparation of matrix spiking

For spiked DNA samples, 1 g of homogenized canned pork or canned tuna (used as the food matrix) was aseptically weighed and transferred into a sterile tube containing 1 mL of bacterial suspension (only for *C. botulinum* serotypes A and B), and 9 mL of sterile distilled water. The bacterial suspensions were prepared by serial ten-fold dilutions in sterile distilled water to achieve final theoretical spiking levels of 10^3^–10^8^ CFU/g in the matrix. For serotypes C through G, the corresponding BoNT encoding gene containing plasmids were used instead of bacterial cells due to the unavailability of strains. The mixture was thoroughly homogenized, and 1 mL of the spiked suspension was collected for DNA extraction.

### DNA Isolation and Quantification

Plasmid DNA was purified using the DNA-spin™ Plasmid DNA Purification Kit (iNtRON, Republic of Korea) and total DNA was extracted using GeneJET DNA Purification Kit (Thermo Fisher Scientific, United States) according to the manufacturer’s instructions. DNA concentration and purity were determined by measuring absorbance at 260 nm (A_260_) using a NanoDrop spectrophotometer. DNA samples with an A_260_/A_280_ ratio between 1.8 and 2.0 were considered sufficiently pure for downstream applications.

The copy number of plasmid DNA templates was calculated using the following formula:

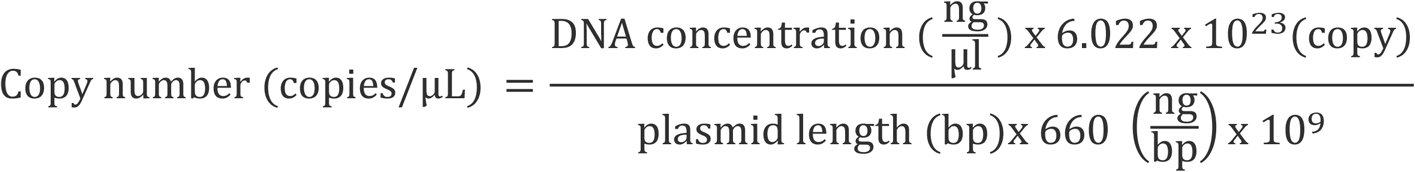

Where 6.022 × 10²³ is Avogadro’s constant, 660 g/mol (or 660 ng/bp) x 10^9^ is the average molecular weight of one base pair (bp), and plasmid length represents the total length of the vector plus the inserted gene fragment (in bp).

### Primer Design and PCR Amplification of *bont/A-G* Genes

Primers specific for amplification of the bont/A gene of *Clostridium botulinum* (Table 1) were designed using PrimerQuest™ (IDT, United States) and Primer-BLAST (NCBI, United States). The sequences of primers were selected to target the conserved region of the genes and avoid maximally the most updated polymorphic nucleotide positions, in some cases degenerate nucleotides were used to cover unavoidable polymorphic nucleotide positions.

Conventional PCR reactions were performed using GoTaq® Green Master Mix. Each 20 µL reaction contained 10 µL of 2× Master Mix, 1 µL of forward primer (10 pmol), 1 µL of reverse primer (10 pmol), 1 µL of template DNA, and 7 µL of nuclease-free water. Negative control reactions contained nuclease-free water instead of template DNA.

PCR amplification was carried out under the following conditions: initial incubation at 95°C for 4 min as a hot start, followed by 40 cycles of denaturation at 95°C for 20 s and primer annealing/extension at 56–62°C for 60 s, depending on primer optimization results. PCR products were analyzed by agarose or polyacrylamide gel electrophoresis.

### Real-Time PCR for Detection and Quantification of BoNT/A–G Genes

Real-time PCR assays were performed using SYBR Green–based master mix (Promega, Thermo Fisher Scientific, or New England Biolabs). Each 20 µL reaction consisted of 10 µL of 2× Master Mix (including Taq/fast DNA polymerase, dNTPs plus dUTP, Uracil-N-glycosylase (UDG), and buffer), 1 µL of forward primer (10 pmol), 1 µL of reverse primer (10 pmol), 1 µL of DNA template, and 7 µL of nuclease-free water. Thermal cycling conditions included an initial incubation at 25°C (or at the optimal temperature of UDG used) for 7 min, followed by a hot start and UDG inactivation at 95°C for 1 min, and 40 cycles of denaturation at 95°C for 15 s and annealing plus extension at 58°C for 1 min. Amplification data were analyzed based on cycle threshold (C_t_) values, amplification curves, and melting curve analysis. Negative controls without template DNA were included in all runs. Selected amplicons were further verified by gel electrophoresis when required. Samples were counted positive if the C_t_ value was less than 40 cycles and its melting curve temperature matched with that of the standard.

### Polyacrylamide Gel Electrophoresis

For separation of small DNA fragments (<1 kb), 8% polyacrylamide gels were prepared in 1× TAE buffer supplemented with ammonium persulfate (75 µL per 10 mL gel) and TEMED (7 µL per 10 mL gel). Samples were mixed with 6× loading dye at a ratio of 5:1 (v/v) and electrophoresed at 120 V for 40–50 min at room temperature. Gels were stained with ethidium bromide (0.1 µg/mL) and visualized under UV light using a GelDoc imaging system.

Specificity of *C. botulinum* BoNT/A-G detection procedure was determined by using the corresponding target gene containing plasmid as the standard at 10^1^-10^7^ copies/reaction. The samples included negative control (without DNA template), the gDNA in case of *C. botulinum* BoNT/A or B or target gene containing plasmid in case of *C. botulinum* BoNT/C-G alone or mixed with other six DNA templates, gDNA of *E. coli* and gDNA of *B. subtillis*. The concentration of each DNA sample was 10^5^ copies/reaction. In addition, the specificity of *C. botulinum* BoNT/A-G detection procedure was determined by using the same DNA templates spiked in canned pork extract. True negative samples are defined as non-target gene containing templates with no proper amplification signal and false positive samples are defined as non-target gene containing templates with proper amplification signal.

Sensitivity of *C. botulinum* BoNT/A-G detection procedure was determined the same way as for specificity, but the concentration of target DNA template was in the range of 5x10^1^-10^7^ copies/reaction. gDNA in case of *C. botulinum* BoNT/A or B or target gene containing plasmid in case of *C. botulinum* BoNT/C-G were spiked in canned pork, DNA was isolated, diluted with 10 mM Tris-HCl buffer containing 2 mM EDTA and used for PCR analysis. True positive samples are defined as target gene containing templates with proper amplification signal and false negative samples are defined as target gene containing templates with no proper amplification signal.

Limit of detection *C. botulinum* BoNT/A-G detection procedure was determined by using the corresponding target gene containing plasmid in the range of 1-10^7^ copies/reaction. The samples included negative control (without DNA template).

### Methodological Rigor

All experiments were performed with appropriate positive and negative controls. Each assay was repeated independently to ensure reproducibility.

## Results

### Identification of the conserved regions and designing of primers for BoNT/A-G encoding genes

To specifically detect all the seven subtypes (A-G) of BoNT genes under the same PCR conditions and to avoid missing the same gene carrying strains due to polymorphism, multiple sequence alignments of the target genes were performed, and the seven most conserved regions (each for one serotype) were selected (Table 1). The length of primers was also selected to have similar melting temperature 61°C/62°C and amplification fragment sizes ranging from 89 bp to 166 bp. Unavoidably, some positions in the selected region for primer design (of BoNT/A, B, E and F) were polymorphic, and degenerate nucleotides were used to cover all possible strains. The specific recognition of each target BoNT gene was checked with *in silico* PCR using online software (insilico.ehu.eus) and only single band of the expected size was obtained for each corresponding *Clostridium botulinum* BoNT serotype (data not shown).

Notably, the positions of the highly conserved regions were different among seven serotypes, with the majority located in the first half of the full-length protein that contains the catalytic endopeptidase domain. There was only one exception, which was for the BoNT/C encoding gene, with the selected region residing in the heavy chain, which interacts with the receptor and is responsible for the translocation of the active enzyme.

### Selection of thermocycle conditions for real-time PCR

Different PCR conditions, including time and temperature for denaturation, primer annealing and extension were tested. As illustrated in Fig.1, for all conditions, only a single amplified DNA band with the expected size of corresponding BoNT genes were obtained. Temperatures 57°C or 58°C were shown to be the best for amplification of all the target BoNT genes (Fig 1). To avoid nonspecific amplification due to primer annealing at low temperature and to eliminate carryover PCR products, commercial master mixes containing UDG and dUTP were selected for real-time PCR experiments. Three master mixes including TaqPath™ 1-Step Multiplex Master Mix (4×; Thermo Fisher Scientific, United States), PowerUp™ SYBR Green Master Mix (Thermo Fisher Scientific, United States), and Luna® Universal qPCR Master Mix (New England Biolabs, United States) showed no considerable difference in specificity and sensitivity (data not shown). We selected an optimal thermocycle conditions, which included 25°C for 7 min for elimination of carryover PCR products by UDG, followed by a hot start at 95°C for 1 min, 40 cycles of denaturation at 95°C for 15 s and annealing plus extension at 58°C for 1 min, the total time of a PCR run was approximately 1.5 hours. Using BoNT/A-G containing plasmids at different concentrations as the standard, the results of real-time PCR assays targeting BoNT/A–G genes produced clear amplification curves with no detectable signal in no-template controls (Fig 2). Mean C_t_ values decreased proportionally with increasing template concentration (logarithm of gene copy number). Standard curves showed that strong linearity with R² values between 0.965 and 0.999. Amplification efficiencies ranged from 100.2% to 109%, with slopes between −3.128 and −3.316 falling within acceptable ranges for quantitative PCR. In addition, each target gene at different concentrations had only one sharp melting curve peak with the same melting temperature.

**Fig 1.**
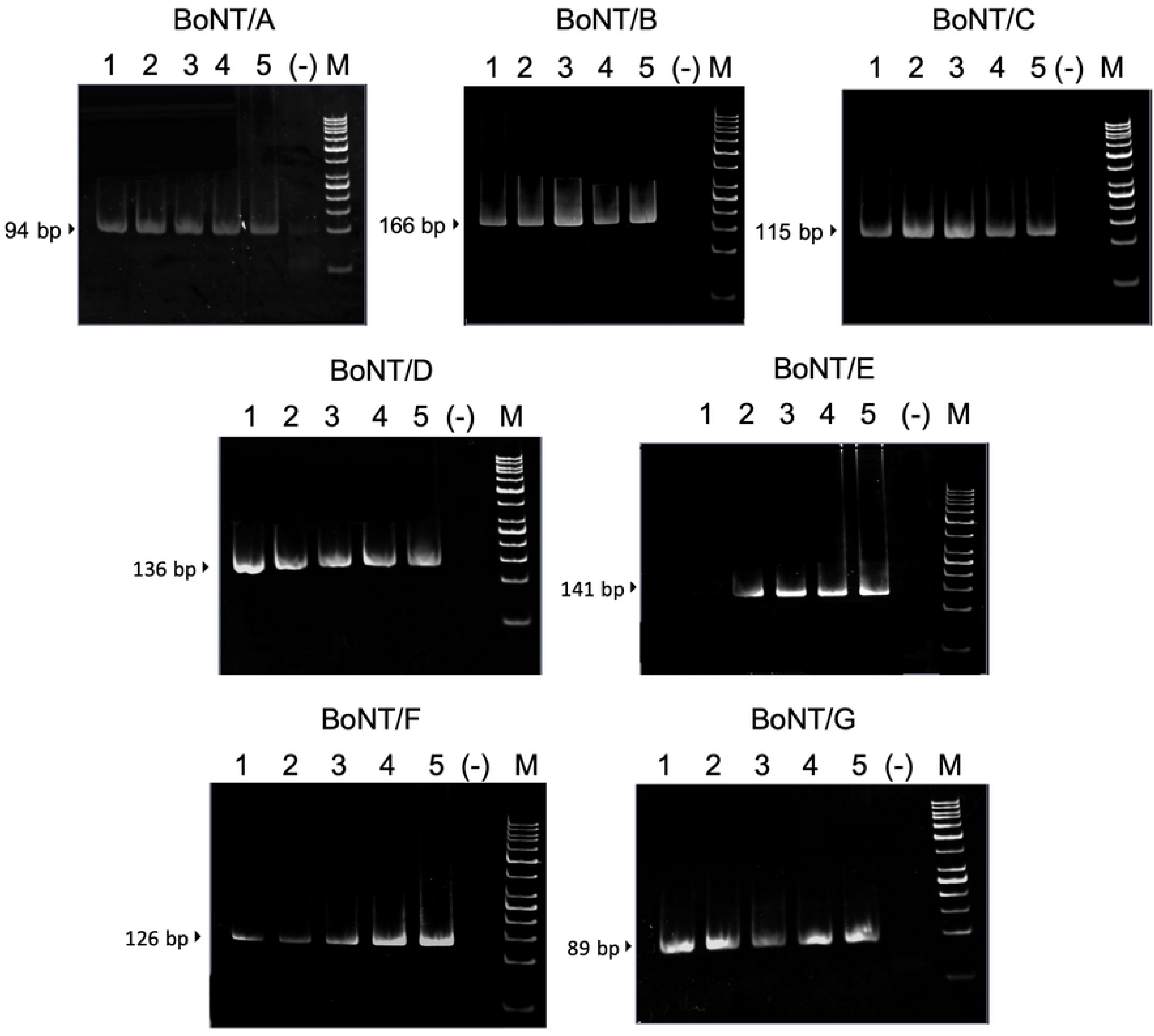
Polyacrylamide gel electrophoresis of BoNT/A-G amplification by PCR under different annelning plus extension temperatures. Lanes 1-7 are for annealing plus extension at 61°C, 60°C, 59°C, 58°C, 57°C, respectively. (-): negative control (no DNA template), M: DNA markers.

**Fig 2.**
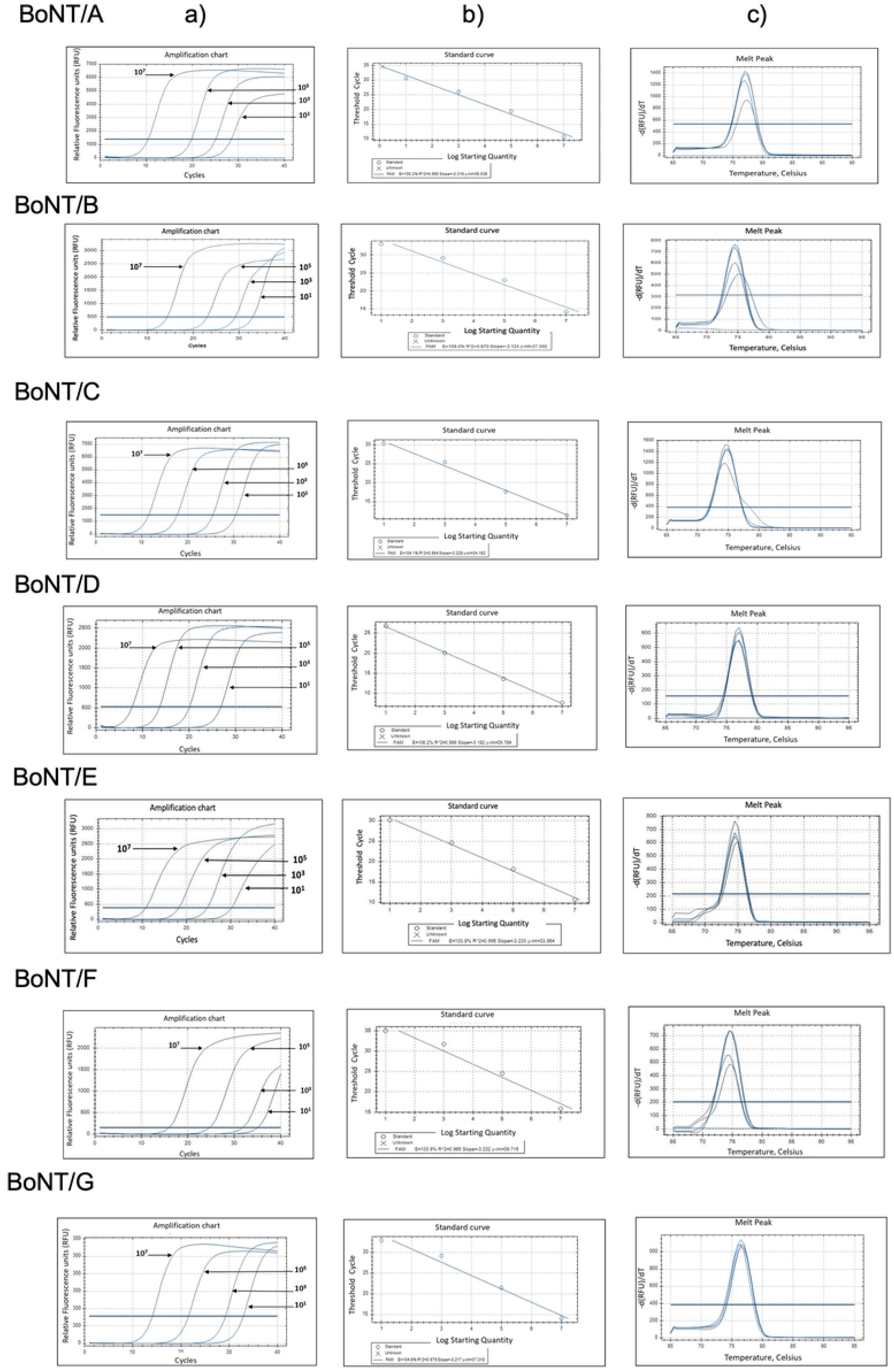
Standard curve construction using control plasmids for BoNT/A-G encoding genes. A) Amplification curves from left to right of BoNT genes at 10^7^, 10^5^, 10^3^ and 10^1^ copies/reaction, respectively, b) The standard curve showing the interrelationship between the threshold (C_t_) values and logarithm of gene copy number, c) The melting curves of PCR products.

### Specificity of the real-time PCR procedure

As shown in Fig 3, using corresponding BoNT gene carrying plasmid alone or mixed with other six BoNT gene carrying plasmids as DNA template, only one amplification curve was obtained with the same sharp melting curve. No amplification signal appeared in other DNA templates including six remaining BoNT genes, gDNA of *E. coli* and gDNA of *B. subtillis*.

**Fig 3.**
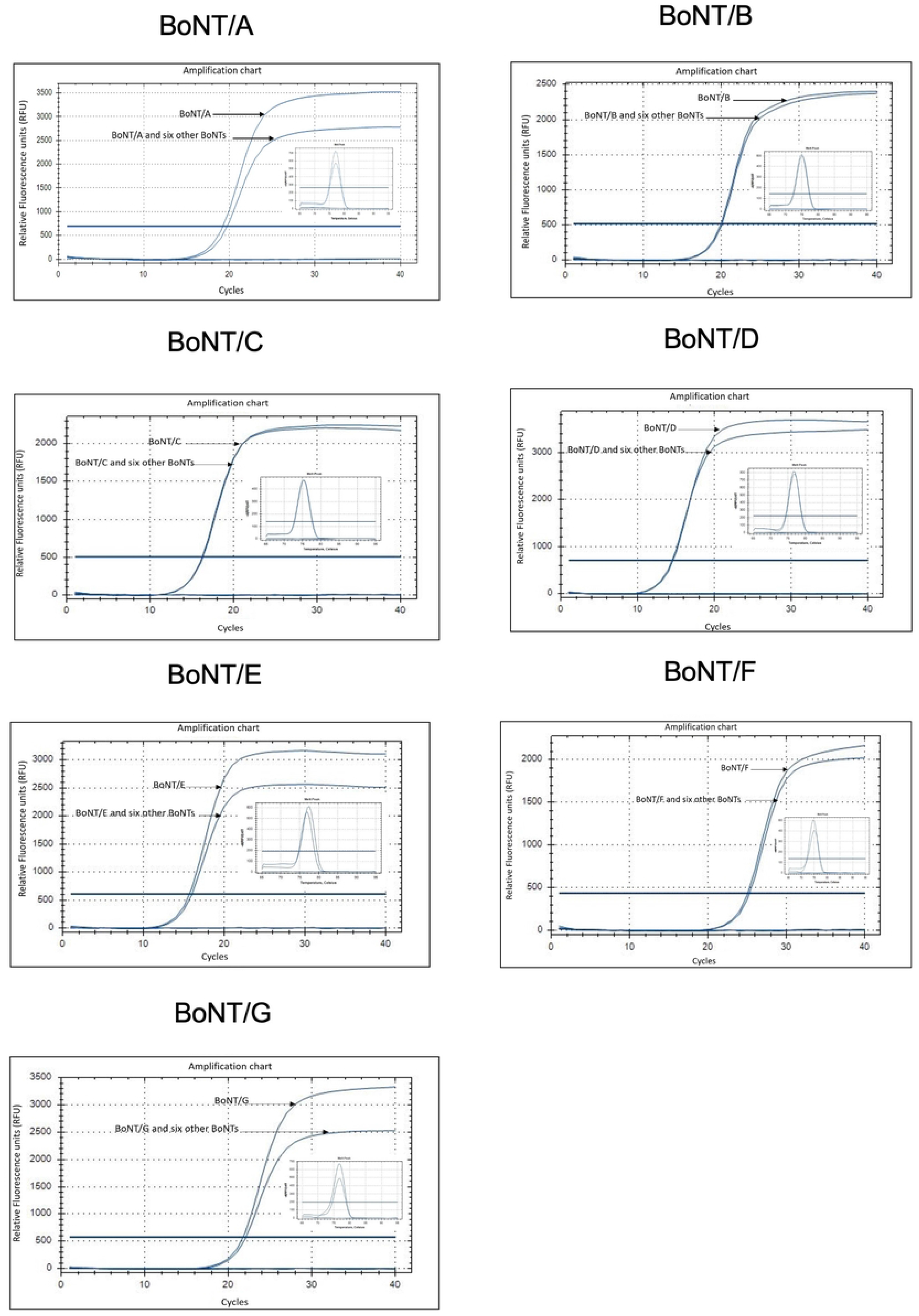
Specificity of real-time PCR procedure in detection of *C. botulinum* BoNT/A-G genes. The amplification curves of BoNT/A-G genes from the different DNA templates and the melting curves of the real-time PCR products (the inset). Reaction mixtures included: No DNA template (negative control), BoNT encoding plasmid of corresponding serotype and BoNT encoding plasmids of 7 serotypes at 10^5^ copies/reaction for each, gDNA of *E. coli* and gDNA of *B. subtillis*.

In other experiments, DNA templates were spiked in canned pork and canned tuna, isolated and used for real-time PCR analysis. The results showed that amplification signal appeared only in corresponding target gene samples (one sample for each target gene) as in the standards, but not in the negative control or the other samples including six remaining BoNT genes, gDNA of *E. coli* and gDNA of *B. subtillis* (Table 2), indicating high specificity of the real-time procedure.

**Table 2.**
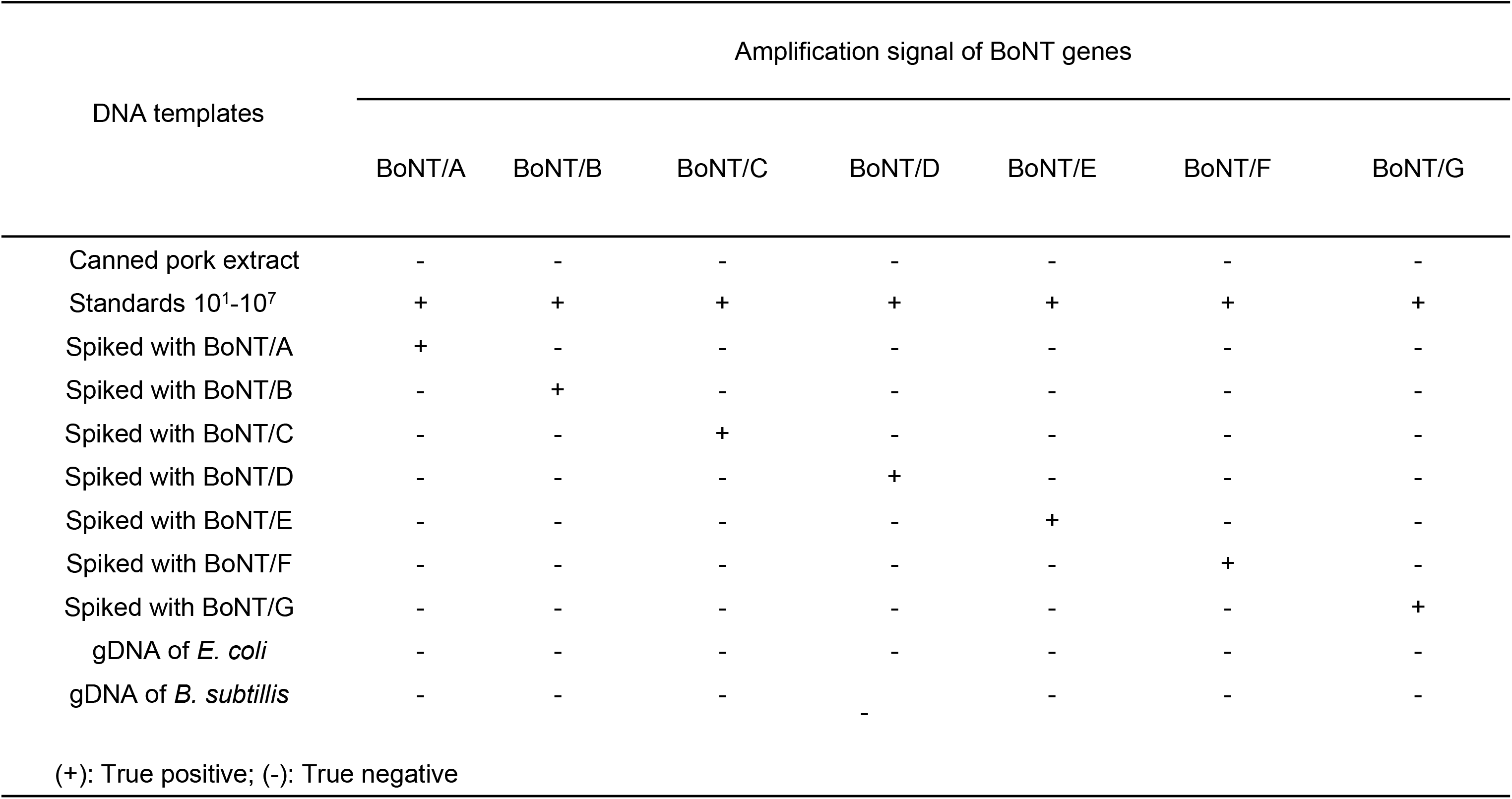
Specificity of real-time PCR procedure for *C. botulinum* BoNT genes.

### Limit of Detection and sensitivity of real-time PCR procedure

In order to determine the limit of detection (LOD), defined as the lowest concentration detected in ≥ 95% of replicates, serial tenfold dilutions containing 10° (1 copy) to 10⁷ copies per reaction of corresponding BoNT gene carrying plasmid were tested. We found that 5 and 10 copies per reaction were the lowest concentrations for PCR detection for BoNT/A-E, G, and BoNT/F, respectively (Table 3).

**Table 3.** Limit of detection of real-time PCR procedure for *C. botulinum* BoNT genes.

| Copy<br>number/reaction | Amplification signal of BoNT genes |  |  |  |  |  |  |
| --- | --- | --- | --- | --- | --- | --- | --- |
|  | BoNT/A | BoNT/B | BoNT/C | BoNT/D | BoNT/E | BoNT/F | BoNT/G |
| No DNA template | - | - | - | - | - | - | - |
| 10<sup>7</sup> | + | + | + | + | + | + | + |
| 10<sup>5</sup> | + | + | + | + | + | + | + |
| 10<sup>3</sup> | + | + | + | + | + | + | + |
| 10<sup>1</sup> | + | + | + | + | + | + | + |
| 5 | + | + | + | + | + | - | + |
| 1 | - | - | - | - | - | - | - |

(+): Positive, (-): Negative

In other experiments, DNA templates were spiked in canned pork, isolated and diluted in a range of 5x10^1^-10^7^ range and used for real-time PCR analysis. The results showed that amplification signal appeared in all the corresponding target gene containing samples, but not in the negative control (Table 4), indicating high sensitivity of the real-time procedure.

**Table 4.**
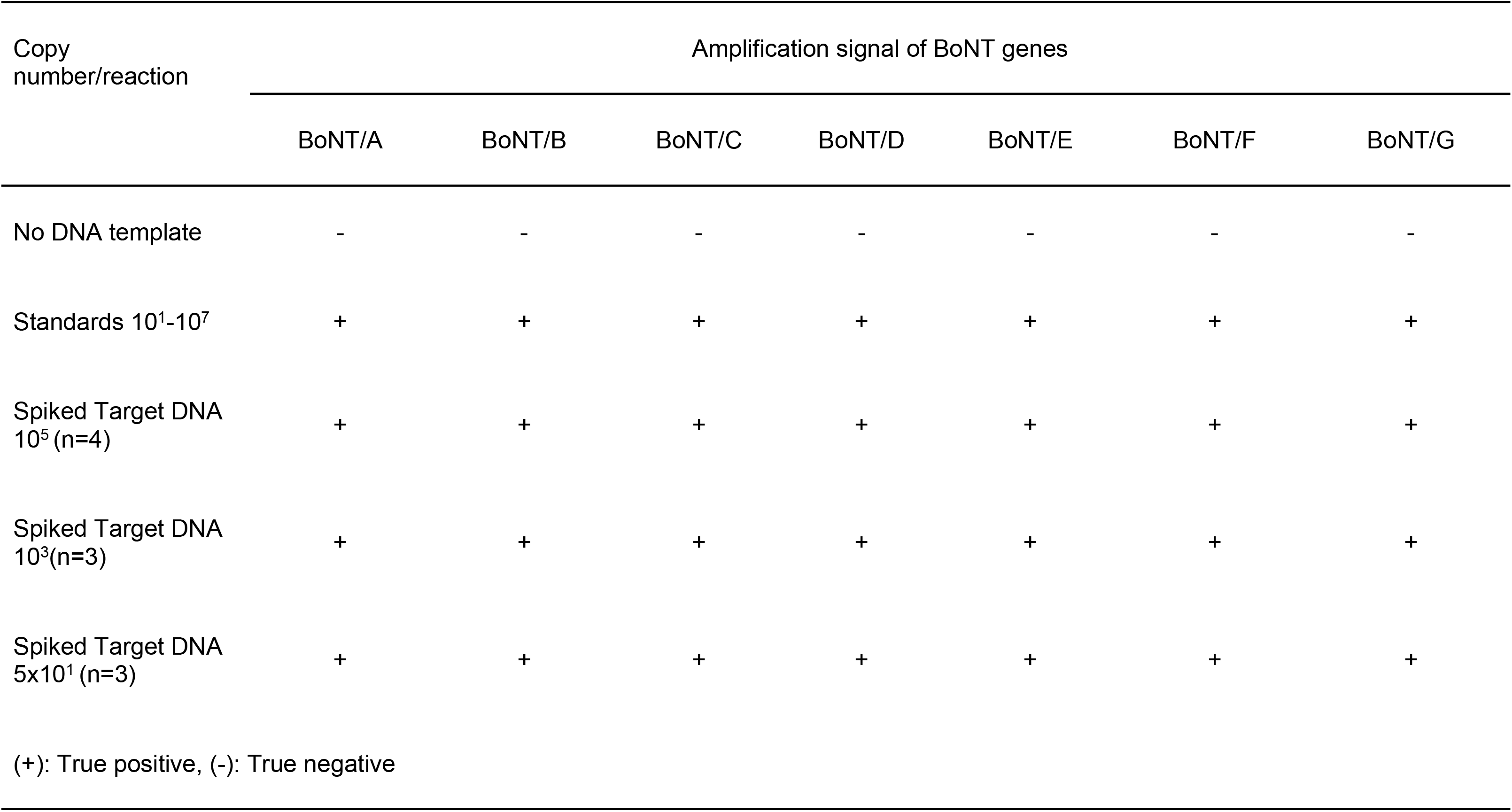
Sensitivity of real-time PCR procedure for *C. botulinum* BoNT genes.

## Discussion

PCR-and real-time PCR-based assays are among the most widely used approaches for determination of the presence of BoNT-encoding genes of *Clostridium botulinum* in botulism investigation because of their versatility and high sensitivity [14, 15–17, 19]. Human botulism is mainly caused by BoNT serotypes A, B, E, and F (less frequent), whereas serotypes C, D, and G are more commonly associated with animal botulism [11, 12]. Many PCR and real-time PCR assays frequently focused on only one or several clinically dominant serotypes, particularly BoNT/A, B, E, and F, rather than comprehensive detection of all toxin groups [14,15–17]. Such approaches may reduce diagnostic coverage in veterinary, environmental, or emerging outbreak settings involving less common serotypes.

The extensive genetic diversity and polymorphism of bont genes remain one of the greatest challenges in molecular detection of *Clostridium botulinum*, because polymorphic positions create mismatches in primer-or probe-binding regions and can markedly reduce amplification efficiency, compromise assay sensitivity, leading to false-negative results [19, 24]. To overcome this limitation, incorporation of degenerate nucleotides in primers were used [15, 17, 19]. In our study, we performed multiple sequence alignments to cover diverse BoNT serotypes and subtypes and identified the most conserved regions for primer design. Even so, in these regions, there existed still some polymorphic positions, and we incorporated degenerate nucleotides in the primers of BoNT/A, B, E and F to compensate for unavoidable polymorphic positions in PCR assay (Table 1). By selecting proper sequence of primers for all the seven BoNT genes, validating primers by *in silico* and conventional PCR, we were able to set up suitable PCR method for detection and quantification of all seven *C. botulinum* BoNT encoding genes under the same conditions.

Our developed real-time PCR assays also demonstrated high analytical performance, with strong linearity between logarithm of gene copy number and C_t_ values (R² values between 0.965 and 0.999). The limits of detection of 5 copies per reaction for BoNT/A-E and G, and 10 copies per reaction for BoNT/F (Table 3) indicate high analytical sensitivity comparable to or better than other previously reported assays [16,18, 20]. In specificity testing, no cross-reactivity was observed with non-target bacterial DNA, including *E. coli* and *B. subtilis*, and amplification signals were only detected from the corresponding BoNT targets even in mixed-template experiments (Fig 3). The high sensitivity and specificity further support the robustness and reliability of the assay for molecular diagnostics.

Carry-over contamination is a well-recognized source of false-positive results in PCR assays, particularly in laboratories performing routine high-throughput testing [25, 26]. To eliminate false-positive results, in our real-time PCR assays, we selected commercial master mixes containing UDG and dUTP. This helps eliminate contamination of uracil-containing amplicons generated from previous reactions without affecting native DNA templates, thus substantially improving the assay reliability while minimizing the risk of false-positive detection.

Overall, our developed real-time PCR assay provides a robust platform for rapid and comprehensive detection of BoNT/A–G genes. The combination of the ability to detect all seven toxin serotypes with broad polymorphism coverage, high sensitivity, specificity, shortened assay time, and effective prevention of carry-over contamination makes the assay suitable not only for clinical diagnosis but also for food safety monitoring, veterinary surveillance, environmental screening, and biodefense applications.

## Conclusion

In summary, we successfully developed and validated a real-time PCR assay for the detection and quantification of BoNT/A–G genes. The assay demonstrated high sensitivity and specificity with limit of detection of 5-10 target gene copy per reaction. By using UDG in combination with dUTP in the PCR assay, the false positive phenomenon due to the carryover PCR product was eliminated. The established procedure could be applied in clinical diagnostics, food safety, and environmental surveillance. The method provides a reliable alternative to traditional bioassays while supporting global botulism surveillance efforts.

## Acknowledgements

This study was financially supported by the Vingroup Innovation Foundation with the grant code VINIF.2022.DA00116.

## Author Contributions

Conceptualization: T-N Phan, Y Pham, T-T Nguyen

Data curation: H-L T Nguyen, T-N Phan

Methodology: T-N Phan, Y Pham, P-L Phan, T-T Le, P-A Le, H-A Chu

Investigation: P-L Phan, H-A Chu, T-T Le, P-A Le, M-N T Tran, H-L T Nguyen, T-T Nguyen, T-N Phan, Y. Pham

Formal Analysis:T-N Phan, Y Pham, P-L Phan, H-A Chu, T-T Le, H-L T Nguyen Project administration: Y Pham

Resources: T-N Phan, Y Pham, M-N T Tran Writing – Original Draft: T-N Phan., Y. Pham Writing – Review & Editing: T-N Phan, Y Pham Visualization: P-L Phan, T-T Le, H-A Chu

Soft ware: P-L Phan, T-T Le Supervision: T-N Phan

Validation: P-L Phan, T-T Le, H-A Chu Funding Acquisition: T-N Phan, Y Pham

## Data Availability Statement

All relevant data supporting the findings of this study are available within the manuscript and its supplementary materials. Additional raw datasets generated and analyzed during the current study are available from the corresponding author upon reasonable request.

## Confict of Interests

The authors declare that they have no conflicts of interest.

## Supporting information

**S1.Table. Sequence and location of selected BoNT/A-G gene containing fragments in the plasmids used as DNA templates and standards**

